# The Complex Landscape of Multidrug Resistance in *Zymoseptoria tritici*: MFS1 Promoter Plasticity and beyond

**DOI:** 10.1101/2023.12.27.573052

**Authors:** Elza Neau, Simon Patry-Leclaire, Cécile Lorrain, Nicolas Lapalu, Anaïs Pitarch, Matéa Vencatasamy, Anne-Sophie Walker, Anaïs Lalève, Sabine Fillinger

## Abstract

Multidrug resistance (MDR) in fungal pathogens poses a growing threat to fungicide efficacy and sustainable agriculture. In the wheat pathogen *Zymoseptoria tritici*, MDR is primarily linked to overexpression of the *MFS1* transporter gene, driven by transposable element (TE) insertions in its promoter or their remnants. Here, we provide a comprehensive analysis of *MFS1* promoter polymorphism and its phenotypic impact on MDR across 374 field isolates collected in Europe between 2020 and 2021. We identify six novel structural variants derived from diverse TE types, confirming that the *MFS1* promoter undergoes recurrent and independent insertion events. While previously characterized inserts consistently confer resistance, other variants show limited or variable phenotypic effects. Genetic crosses and quantitative phenotyping further reveal that MDR behaves as a quantitative trait and genome-wide association study confirmed *MFS1* as the major resistance locus but also identified additional candidate genes involved in xenobiotic detoxification and membrane transport, supporting a polygenic basis for MDR. Our findings highlight the interplay between TE-driven structural variation and background polygenic architecture in shaping resistance phenotypes. In parallel, this study highlights the need to improve MDR monitoring beyond single-locus genotyping and re-evaluate current resistance management strategies that may inadvertently select for broad-spectrum resistance.

## Introduction

Major advances in public health and agriculture over the past century were largely driven by the development and widespread use of pesticides and medical drugs, referred to as active ingredients (AIs). These innovations have significantly reduced disease-related mortality and increased crop yields (OECD, 2018; Oerke, 2006). However, the extensive use of AIs imposes strong selective pressure on microbial populations, generally leading to resistance development. This growing concern is not limited to bacterial pathogens. In recent years, fungal resistance to some AIs (*e.g.*, inhibitors of 14α-demethylation) has emerged as a parallel threat, both in clinics and agriculture where its impact on global food security is underestimated (Fisher and Denning, 2023; Stukenbrock and Gurr, 2023). Characterizing fungicide resistance has historically focused on the identification of target-site resistance (TSR), arising from mutations or overexpression of the target protein (R4P Network, 2016). Such mechanisms, often associated with high resistance factors, have been central to resistance management strategies for decades (Lucas *et al*., 2015). To slow down their selection, growers are encouraged to alternate or mix fungicides of different modes of action (MoAs) (Rex Consortium, 2013). Despite the widespread application of fungicides and the deployment of alternative control methods such as resistant crop varieties, global agricultural losses due to fungal pathogens remain significant, estimated between 10% and 23% annually (Fisher *et al*., 2012). Among the causes of these persistent losses, the rise of fungicide-resistant strains is a significant factor, undermining the efficacy of current management strategies. This highlights that advancing our understanding of fungicide resistance is essential to sustain crop productivity and address future food security challenges.

Multidrug resistance (MDR) refers to mutation(s) affecting genes unrelated to the target of the AI (non-target-site mutations) and conferring broad-spectrum resistance to multiple fungicides with unrelated MoAs (R4P Network, 2016). The most commonly described mechanism underlying MDR in fungal plant pathogens involves increased efflux activity typically mediated by membrane transporters of the ATP-binding cassette (ABC) family and major facilitator superfamily (MFS) (Hu and Chen, 2021). MDR has been well documented in human health including in cancer (Krishna and Mayer, 2000), and since the early 2000s, mounting evidence has revealed its presence in phytopathogenic fungi such as *Botrytis cinerea*, *Oculimacula* spp., *Penicillium digitatum*, *P. expansum*, *Clarireedia homoeocarpa* (formerly *Sclerotinia homoeocarpa*), and *Zymoseptoria tritici* (Kretschmer *et al*., 2009; Leroux *et al*., 2013; Leroux and Walker, 2013, Sang and Hulvey, 2018; Sun *et al*., 2013). In the necrotrophic grey mold agent *B. cinerea*, MDR results from two different mutational events that recombined in some isolates. The gain-of-function mutation in the transcription factor BcMrr1 leads to constitutive overexpression of the ABC transporter BcAtrB, while overexpression of the MFS transporter BcMfsM2 is driven by retrotransposon insertion in its promoter (Kretschmer *et al*., 2009). In the turfgrass dollar spot agent *C. homoeocarpa*, MDR is linked to the upregulation of three cytochrome P450 enzymes and two ABC transporters, controlled by a gain-of-function mutation in the transcription factor ShXDR1 (Sang and Hulvey, 2018). These examples highlight the complex genetic architecture of MDR, which can arise from diverse and independent events and may be combined in populations *via* sexual recombination. If MDR *per se* is associated with low to moderate resistance factors (RFs), its potential to both broaden the resistance spectra independently from MoAs and significantly elevate RFs when associated with TSR poses a serious threat to the efficacy of both registered and novel fungicides (Hu and Chen, 2021; Mitchell *et al*., 2014; Vázquez-García *et al*., 2020). Because of this potential to compromise the efficacy of future MoAs, including biocontrol molecules, and given the slow pace of AI discovery (Renwick and Mossialos, 2020; Sparks and Bryant, 2021), MDR surveillance should be a key priority for sustainable resistance management, particularly through the development of field-adapted monitoring tools.

*Zymoseptoria tritici* the causal agent of the Septoria Tritici Blotch (STB) disease, is the most damaging wheat leaf pathogen in temperate regions (Orton *et al*., 2011; Dean *et al*., 2012, Klink *et al*., 2021). Yield losses attributed to this ascomycete fungus vary annually, ranging from 5% to 50% (Fones and Gurr, 2015). In Europe, the control of STB relies heavily on fungicide applications, with an estimated market size of $1.2 billion (Torriani *et al.et al.*2015; Figueroa, Hammond-Kosack, and Solomon 2018) accounting for approximately 70% of annual fungicide usage dedicated to wheat. Several fungicides with different MoAs are used to manage STB in the field, including multisite inhibitors (folpet, sulphur), sterol 14α-demethylation inhibitors (DMIs), succinate dehydrogenase inhibitors (SDHIs), and more recently the quinone inside inhibitor (QiI) fenpicoxamid (Garnault *et al*., 2019). In France, a typical fungicide program includes DMI and/or multisite applications followed by mixtures of DMI and SDHI or QiI, particularly for the critical sprays targeting the emergence of the first leaf (ARVALIS, 2024, 2016). However, *in vitro* experimental evolution studies have revealed that the counterpart of diversifying selection pressure through such mixture or alternation could be the inadvertently emergence of MDR in *Z. tritici,* but also in other organisms (Ballu *et al*., 2024, 2023, 2021; Barbosa *et al*., 2021; Comont *et al*., 2020; Lagator *et al*., 2013; Zhou *et al*., 2022).

MDR in *Z. tritici* was first identified in DMI-resistant field isolates from France, displaying moderate cross-resistance to DMIs, SDHIs, QoIs, and even to squalene epoxidase inhibitors (SBIs of class IV, namely terbinafine, tolnaftate) a MoA not used in agriculture (Leroux and Walker, 2011). This phenotype was linked to the constitutive overexpression of the *MFS1* gene, encoding an MFS transporter (Omrane *et al*., 2015, 2017). In most cases, overexpression was driven by a 519 bp long-terminal repeat (LTR) insertion in the *MFS1* promoter region (*P_MFS1_*), designated as the type I insert (Omrane *et al*., 2017). Additional insertions, types II and III, have also been identified, each contributing to varying levels of *MFS1* expression and MDR phenotypes (Omrane *et al*., 2017). Such transposable elements (TE)-derived insertion polymorphisms exemplify an important role of TEs in driving genome evolution and adaptive responses in *Z. tritici* (Baril *et al*., 2025; Lorrain *et al*., 2021; Möller and Stukenbrock, 2017; Oggenfuss and Croll, 2023; Singh *et al*., 2021). Field monitoring from 2008 to 2017 in France showed a rise and stabilization of MDR frequency at approximately 25%, with ∼75% of MDR isolates carrying the *P_MFS1_* ^type I^ insert (Garnault *et al*., 2019). However, this dominance of *P_MFS1_* ^type I^ insert is not observed across Europe as a whole. In Ireland, contrasting insertion frequencies were reported between field populations from different regions, some dominated by type II and III inserts (Kildea *et al*., 2024). Recently, new insertions, referred to as types IV and V, have been described (Lavrukaitė *et al*., 2023; Mäe *et al*., 2020), but no clear phenotypic association with MDR has been demonstrated. A genotyping approach based on PCR amplicon size upstream of *MFS1* is widely used to diagnose MDR in *Z. tritici*. However the accuracy of this approach is limited by the high polymorphism of the locus (Huf *et al*., 2018; Lavrukaitė *et al*., 2023; Mäe *et al*., 2020; Omrane *et al*., 2015, 2017) and by insertions that are not functionally linked to resistance, such as the type V. Furthermore, MDR phenotypes can emerge without any *P_MFS1_* insertion, as demonstrated by recent experimental evolution studies. Therefore, a diagnostic approach solely based on *P_MFS1_* insertion polymorphism is insufficient for reliably monitoring MDR evolution *in natura*.

The present study provides an in-depth analysis of *P_MFS1_* insertion polymorphisms, and associated terbinafine resistance variation, in a collection of 374 *Z. tritici* isolates sampled between 2020 and 2021. Terbinafine resistance was used as a proxy for MDR based on the cross-resistance to SBIs of class IV, DMI, QoI and SDHI conferred by *P_MFS1_ ^type I^* as demonstrated by Omrane *et al*.(2017). By combining PCR-based screening with whole-genome sequencing, we uncovered an unprecedented level of promoter diversity in the *P_MFS1_* region, significantly expanding the known allelic landscape and revealing where current MDR genotyping protocols may lack resolution. Specifically, we identified six new structural variants within the proximal *P_MFS1_* promoter, not consistently associated with the MDR phenotype. We also showed that *P_MFS1_* polymorphisms are not solely responsible for the MDR phenotype, and uncovered by a genome-wide association study (GWAS) a polygenic basis. Several candidate resistance-related genes were identified.

## Results

### The *MFS1* promoter is subject to TE-Mediated structural variation in multidrug-resistant populations of *Z. tritici*

To unravel the *P_MFS1_* evolution and its correlation with MDR in *Z. tritici*, we assembled a collection of 374 field strains collected from Northern Europe between 2020 and 2021 (Table S1). Potential MDR strains were discriminated with tolnaftate or terbinafine, two squalene epoxidase inhibitors (SBIs of class IV) commonly employed as proxies for detecting MDR in field populations (Leroux & Walker, 2011). We obtained 283 potential MDR isolates selected from the French field populations in these two years. Additional potential MDR isolates were provided by collaborators, including 25 from Germany, 26 from the United Kingdom, 22 from Ireland, 12 from Lithuania, 4 from Belgium, 1 from Estonia and 1 from Denmark (Table S1).

We first genotyped the promoter of the *MFS1* gene (*P_MFS1_*) of all 374 isolates using a PCR-based assay targeting the proximal promoter, 500 bp upstream of the start codon. Based on amplicon size and Sanger sequencing of selected isolates, we identified several *P_MFS1_* genotypes including: type I (1005 bp), type II (750-855 bp), type III (636 bp), and *P_MFS1_^no-insert^* (486 bp, ± 50 bp), as well as multiple amplicons of unexpected sizes (Fig. 1A). Insert types were assigned based on size and sequence similarity to previously characterized elements (Omrane *et al*., 2017). The genotyping results revealed a high level of *P_MFS1_* promoter diversity. As only the French isolates (n=283) were randomly selected, the relative insert proportion should reflect that of French wild MDR populations. The already described insertion *P_MFS1_^type I^* was the most frequent genotype (∼70%, n=195), followed by *P_MFS1_^type II^* (∼12%, n=34 for type IIA and IIB), and *P_MFS1_^type III^* (∼5%, n=15) (Table S1, Fig. 1A). Notably, approximately 10% of isolates selected on terbinafine or tolnaftate from the French collection displayed no detectable insertion, *P_MFS1_^no-insert^* genotype (n=28) (Table S1, Fig. 1A). We detected seven new insertions at the *P_MFS1_* locus. Four new insertions were identified at the *P_MFS1_^type II^* insertion locus (471 bp upstream of the *MFS1* start codon) including *types P_MFS1_^type IIC^, P_MFS1_^type IID^, P_MFS_^type IIE^,* and *P_MFS1_^type IIF^* (sizes 267 bp to 339 bp), which shared conserved domains with the known *P_MFS1_^type IIA^* and *P_MFS1_^type IIB^* variants (Fig. 1A and B). The fifth insert at the *P_MFS1_ ^type II^* site, lacking sequence similarity to the others, was designated *P_MFS1_ ^type VII^*. We identified a sixth novel insert, *P_MFS1_ ^type VI^* found 215 bp upstream of the *MFS1* start codon and with a 360 bp size. Finally, one structural rearrangement consisting of a 787 bp deletion from the proximal promoter (108 bp upstream of the *MFS1* start codon) combined with a 159 bp insertion. This new structural variation was named *P_MFS1_ ^type VIII^.* All these novel inserts were found in French isolates, except *P_MFS1_ ^type IIC^* and *P_MFS1_ ^type^ ^IIE^* that were also detected in strains from Lithuania (n=3) and Germany (n=3). These findings expand the known repertoire of *P_MFS1_* insert polymorphisms and suggest that the *MFS1* promoter region is prone to insertion events.

**Fig. 1:**
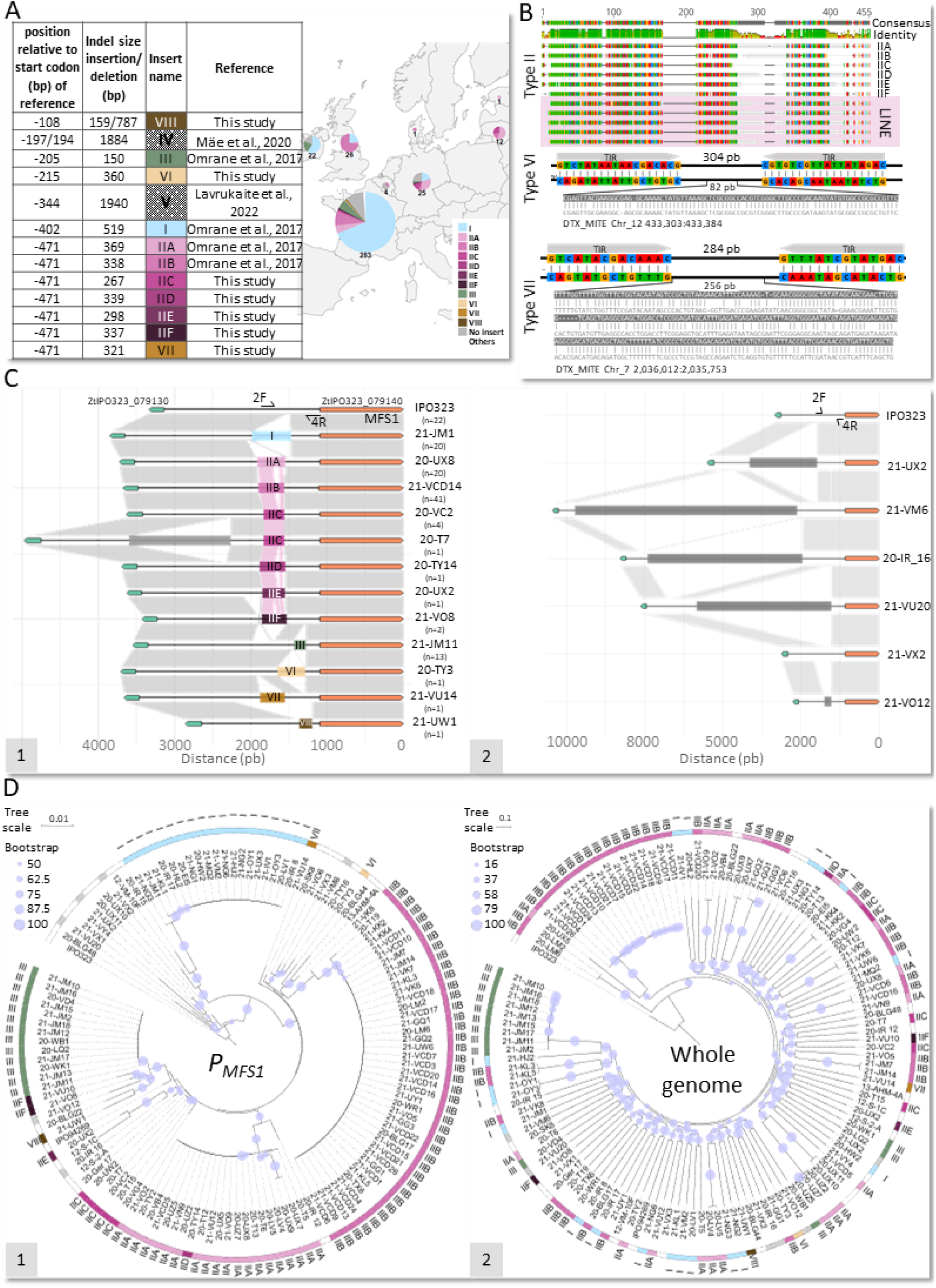
MFS1 promoter (*P_MFS1_*) of *Zymoseptoria tritici* European field strains shows an extended transposon insertion polymorphism. **A. Summary of inserts detected by PCR amplicon sequencing of the proximal *P_MFS1_* and geographic origin of detected polymorphisms from 2020-2021 survey.** Inserts are listed according to their localization relative to the start codon of *MFS1* in the IPO323 reference genome. Insert numbering is according to their chronological discovery. Frequencies shown for French isolates reflect proportions of each *P_MFS1_* genotype in multidrug-resistant (MDR) field populations. For other European countries, strains were provided by partners because they displayed particular *P_MFS1_* amplicon size or sequence. Inserts indicated with a dotted pattern were reported in previous studies but not detected in this survey. “Others” refers to genotypes not characterized by PCR amplicon sequencing due to amplification failure. **B. New *P_MFS1_* inserts found in *Z. tritici* field isolates contain transposable element signatures.** Type II insert variants were aligned with the end of LINE retrotransposons of the IPO323 genome (pink highlighting). Sequence identity is shown by colored bases. Type VI and VII inserts contain terminal inverted repeats (TIRs, grey boxes) and internal sequences aligned to known MITEs from IPO323 as indicated by colored bases. **C. *P_MFS1_* sequence reassembly from whole genome sequencing reads reveals insertion polymorphism upstream of the routinely genotyped proximal promoter by amplification with 2F/4R primer pair.** A subset of 130 isolates from the 2020–2021 MDR survey were sequenced (Illumina, 25× coverage, 150 bp reads). *P_MFS1_* sequences were extracted from genome assemblies, aligned with MUSCLE, and manually curated. Haplotypes (based on indels >100 bp) are shown with a representative strain’s name and the number of strains sharing that haplotype (n). 2F/4R primer hybridization sites (Omrane et al., 2015) on the IPO323 reference are indicated. C1: haplotypes with insertions detected within the proximal *P_MFS1_ via* 2F/4R PCR amplification. C2: haplotypes harboring insertions (dark grey boxes) not captured by 2F/4R PCR genotyping. Lengths of uncharacterized indels are illustrative, as genome reassembly is unreliable for repetitive elements. Conserved sequences relative to IPO323 are shown as grey polygons, shared sequences among type II inserts are shown in pink. **D. Phylogenetic analysis indicates local conservation of *P_MFS1_*.** Maximum likelihood phylogenies were inferred with IQ-TREE from aligned *P_MFS1_* 2,074 positions shared by all strains (D1) or synthetic sequences made of variants distributed evenly across the genome. Node supports were assessed with 1000 bootstrap replicates (percentages shown). Colors indicate insert types, consistent with panel A. Branch lengths are proportional to the expected number of substitutions per site, indicated by the tree scale.

To explore the origin of these insertions, we aligned their sequences against the IPO323 reference genome, and performed similarity searches to IPO323 annotated transposable elements (Baril & Croll, 2023). This analysis confirmed that nearly all novel and known insertions are derived from or share homology with known TEs, including LTR retrotransposon (*P_MFS1_ ^type I^*), Long interspersed nuclear element (LINE) retrotransposons (*P_MFS1_ ^type IIA^, P_MFS1_ ^type IIB^, P_MFS1_ ^type IIC^, P_MFS1_ ^type IID^, P_MFS1_ ^type IIE^, P_MFS1_ ^type IIF^*), unclassified TE (*P_MFS1_ ^type III^*) and miniature inverted-repeat transposable elements (MITEs) (*P_MFS1_ ^type VI^*, *P_MFS1_ ^type VII^*), Fig. 1B. Finally, the *P_MFS1_ ^type VIII^* variant did not match any annotated TE in the IPO323 genome. However, the inserted fragment displayed partial homology (40-55%) to repetitive sequences on chromosomes 3, 5, and 11, suggesting it may originate from a high copy element (data not shown). These results indicate that recurrent transposition events shaped the structural polymorphism of the *MFS1* promoter region (Fig. 1C-1), establishing this locus as a potential hotspot for TE insertions among MDR strains able to grow on SBI-class IV fungicides.

To further investigate the evolutionary history and origin of *P_MFS1_* insertions, we conducted a phylogenetic analysis of the *MFS1* promoter region based on *de novo* assemblies of 131 strains. We first aligned the 3 kb intergenic region between *ZtIPO323_079130* and *ZtIPO323_079140* (*MFS1,* Fig. 1C) of the reference isolate to examine large-scale structural variation beyond the proximal promoter. This revealed previously undetected structural variations even in three strains initially genotyped as *P_MFS1_*^no-insert^, as well as three strains we failed to amplify by PCR (Fig. 1C-2). Strain 21-VU20 carries a large insertion of ∼5 kb, while strains 21-VO12 and 21-VX2 exhibit large deletions of 757 bp and 322 bp, respectively. These structural variants, often located around the 401 bp upstream relative to the *MFS1* start codon, underline the structural plasticity of the *MFS1* promoter and confirm that current genotyping approaches may overlook substantial structural variant diversity.

**Fig. 2:**
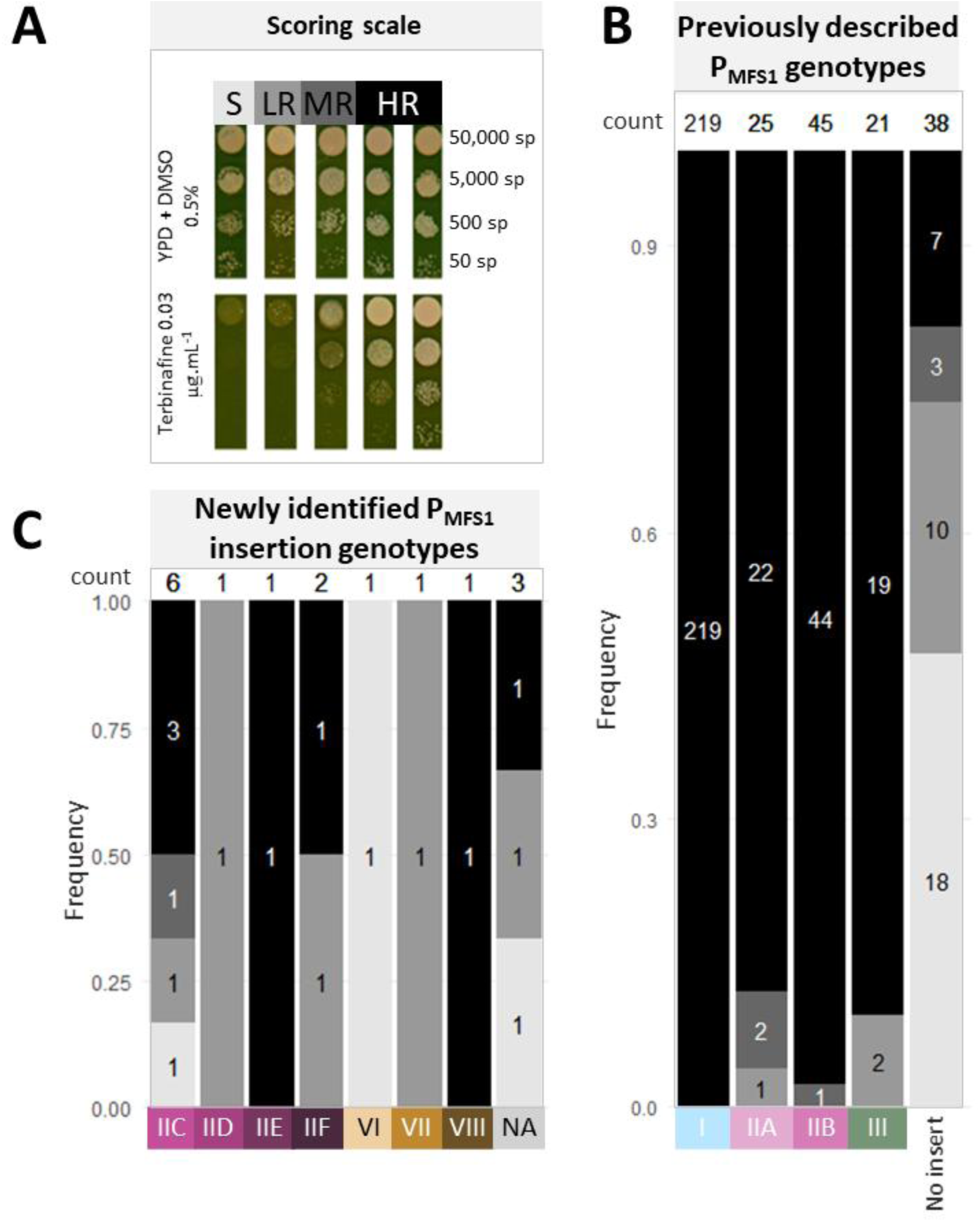
Qualitative terbinafine resistance assessment reveals intra and inter *_PMFS1_* genotype phenotype variations. A total of 374 strains selected on terbinafine or tolnaftate from the 2020–2021 collection was genotyped for insertions in the 500-pb proximal ***_PMFS1_*** sequence *via* PCR amplicon sequencing. The strains were also screened for terbinafine resistance through growth tests on YPD solid medium supplemented with discriminatory terbinafine dose (0.03 µg.mL^-1^ A. scoring scale used to categorize strains Strains are grouped by insertions: **B. previously associated with MDR (I, IIA, IIB, IIC)** and **C. newly identified insertions (IID, IIE, IIF, VI, VII, VIII) as well as NA for uncharacterized genotypes** due to PCR amplification failure. Resistance classes: HR = high resistance, MR = medium resistance, LR = low resistance, S = sensitive.

To better understand the evolutionary dynamics underlying this structural diversity, we next examined the sequence conservation and divergence of the *P_MFS1_* region through phylogenetic analysis. We constructed a maximum likelihood tree based on a multiple sequence alignment (MSA) of 2,074 positions shared in *P_MFS1_* across the 131 strains, excluding insert sequences. This revealed that promoter sequences carrying the same insert, particularly *P_MFS1_ ^type I^* and *P_MFS1_ ^type III^*, formed well-supported monophyletic clades, indicating strong conservation of surrounding promoter regions. Similarly, *P_MFS1_* sequences carrying the same *P_MFS1_ ^type II^* insert variants (IIA, IIB, IIC, IID, IIE and IIF) showed high internal sequence similarity, but did not cluster together across subtypes (Fig. 1D-1). These findings argue against the hypothesis of a single ancestral *P_MFS1_ ^type II^* insertion that diversified through mutation and recombination. Instead, they suggest that the different *P_MFS1_ ^type II^* variants likely arose from multiple, independent transposition events. One exception may be the *P_MFS1_ ^type IID^* variant, which clustered closely with *P_MFS1_ ^type IIA^*, possibly reflecting a shared origin. To assess whether these patterns were driven by population structure, we compared the *P_MFS1_*-based tree to a whole-genome SNP phylogeny (Fig. 1D-2). While some clonal groups were apparent (e.g., strains 21-VCD and 21-JM), the broader patterns of *P_MFS1_* conservation could not be explained by population structure alone. Finally, we quantified nucleotide diversity (π) across PMFS1 haplotypes, retaining a single representative per clonal group. *P_MFS1_* haplotype of strains lacking insertions (*P_MFS1_^no-insert^*) showed tenfold to fiftyfold higher diversity (π = 0.0197, n=13) than insert-carrying haplotypes such as *P_MFS1_^type I^* (π = 0.0004, n=18), *P_MFS1_ ^type IIA^* (π = 0.0017, n=14), *P_MFS1_ ^type^ ^IIB^*(π = 0.001, n=18), and *P_MFS1_ ^type III^* (π = 0, n=5). These patterns are consistent with a local selection of this locus, maintaining specific insertion haplotypes.

### *P_MFS 1_* insertion variants quantitatively impact multidrug resistance in *Z. tritici*

To dissect the contribution of structural polymorphisms at the *MFS1* promoter to MDR in *Z. tritici*, we evaluated terbinafine resistance across field isolates grouped by *P_MFS1_* genotype, as a proxy for MDR characterization. Both qualitative resistance profiling (Fig. 2) and quantitative EC₅₀ measurements were performed (Fig. 3), allowing a high-resolution assessment of phenotypic outcomes associated with individual insertions (Fig. 3A-1). For qualitative resistance profiling, strains were assigned to three terbinafine resistance classes according to their growth score on terbinafine supplemented media (growth scale in Fig. 2A), namely Low Resistance (LR), Medium Resistance (MR) and High Resistance (HR). Strains carrying the *P_MFS1_^type I^* insertion are consistently associated with HR (Fig. 2B). EC₅₀-based phenotyping of 19 randomly selected strains confirmed the strongest resistance, with field isolates displaying resistance factors (RFs) ranging from 2.5 to 15.4 compared to the sensitive IPO323 reference (Fig. 3A-3). The *P_MFS1_^type I^* reference strain Ref_I, which carries the *P_MFS1_^type I^* insertion in the sensitive IPO323 background (Omrane *et al*., 2017), was included in all replicates as a positive control. Most *P_MFS1_^type I^* field isolates showed EC₅₀ values comparable to or exceeding Ref_I (RF 5.1), with some strains such as 21-JM1 reaching an RF of 15 (Fig. 3A-3). These results confirm that *P_MFS1_^type I^* insertions robustly contribute to the MDR phenotype in *Z. tritici*.

**Fig. 3.**
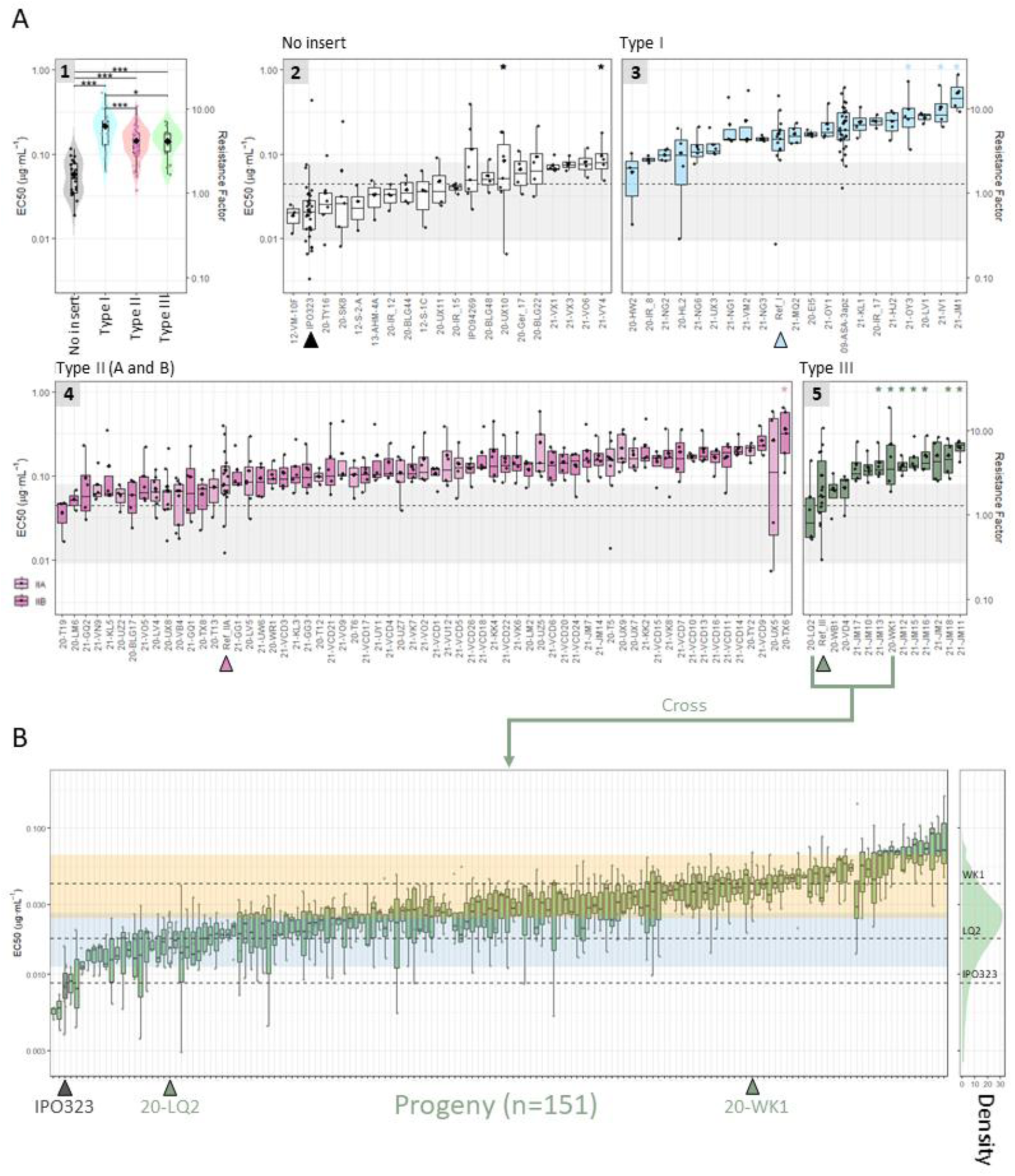
Distribution of terbinafine resistance demonstrates the quantitative nature of MDR. A. Variation of terbinafine EC₅₀ among MDR field strains. (1) Violin plot showing the effect of P*_MFS1_* genotype on terbinafine EC₅₀ (EC₅₀ distributions per genotype). Each violin represents the distribution of EC₅₀ values for strains grouped by P*_MFS1_* genotype. Each point corresponds to the mean EC₅₀ of a strain (biological replicates (n ≥ 2), and the larger points represent the mean of the P*_MFS1_* group. Genotype effects were assessed using a linear mixed-effects model on log-transformed EC₅₀, including biological variation and strain as random effects. Estimated marginal means were compared with contrast post-hoc tests and Bonferroni correction (* *P* < 0.05; *** *P* < 0.001). (2-5) Boxplots of terbinafine EC₅₀ per strain grouped by genotypes, showing intra-genotype variability independent of *P_MFS1_* effect, including experimental variation. The experimental variability of biological replicates (n ≥ 2) is shown by individual points and error bars (standard deviation of biological replicates). Diamond-shaped points indicate the mean EC₅₀ per strain. Each group displays a reference standard, highlighted by colored arrows, carrying the corresponding P*_MFS1_* genotype, *i.e*., Ref_I, Ref_II and Ref_III are IPO323 transformants with *in locus* type I, IIA, or III insertions. Statistical differences with the corresponding reference strain were assessed using a linear mixed-effects model with post-hoc contrasts and Bonferroni correction (* *P* < 0.05). Resistance factor (right y-axis) represents the ratio of a strain’s mean EC₅₀ relative to sensitive control IPO323. **B. terbinafine EC₅₀ in progeny from 20-LQ2 × 20-WK1 type III cross shows a quantitative distribution.** EC₅₀ values of individual progeny derived from sexual crosses between type III *Pmfs1* field strains 20-LQ2 and 20-WK1 follow a normal distribution. Individual points represent biological replicates (n ≥ 2). Boxplots of parental strains are pointed out with green arrows and IPO323 susceptible control with grey one. Dotted lines indicate mean EC₅₀ of IPO323, 20-LQ2, and 20-WK1. Orange and blue boxes indicate the IC95 boundaries of EC₅₀ for 20-WK1 and 20-LQ2, respectively. Density curve of mean EC₅₀ values is shown on the right, summarizing the distribution of mean EC₅₀ values across all progeny. For both A and B, EC₅₀ determination, growth in liquid YPD medium supplemented with increasing terbinafine concentrations was monitored at OD405_nm_ after 7 days (18°C, dark) and expressed as percentage of growth relative to control. EC₅₀ was computed from dose-response curves fitted to each technical replicate. All EC₅₀ values are represented log10-transformed for clarity.

Similar trends were observed for *P_MFS1_^type II^* insertions. Qualitatively, both *P_MFS1_^type IIA^* and *P_MFS1_^type IIB^* were strongly associated with MDR, with 88% (22/25) of *P_MFS1_^type IIA^* and 97% (44/45) of *P_MFS1_^type IIB^* strains of HR phenotype (Fig2B). EC₅₀ quantification supported this association. Field strains carrying *P_MFS1_^type IIA^* or *P_MFS1_^type IIB^* showed significantly higher terbinafine EC₅₀ values compared to IPO323 (Fig. 3A-4) with RFs ranging between 2 and 10. The *P_MFS1_^type II^* reference strain Ref_IIA, carrying a type IIA insert in the IPO323 background, served as the positive control for this group. Most field strains were comparable to Ref_IIA (RF 3.4), with only 20-TX6 displaying significantly threefold higher EC_50_ mean. As for *P_MFS1_^type I^*, this demonstrates that *P_MFS1_^type IIA^* and *P_MFS1_^type IIB^* insertions are functionally linked to the highest MDR levels in the collection, although some inter-strain variability was observed.

**Fig. 4:**
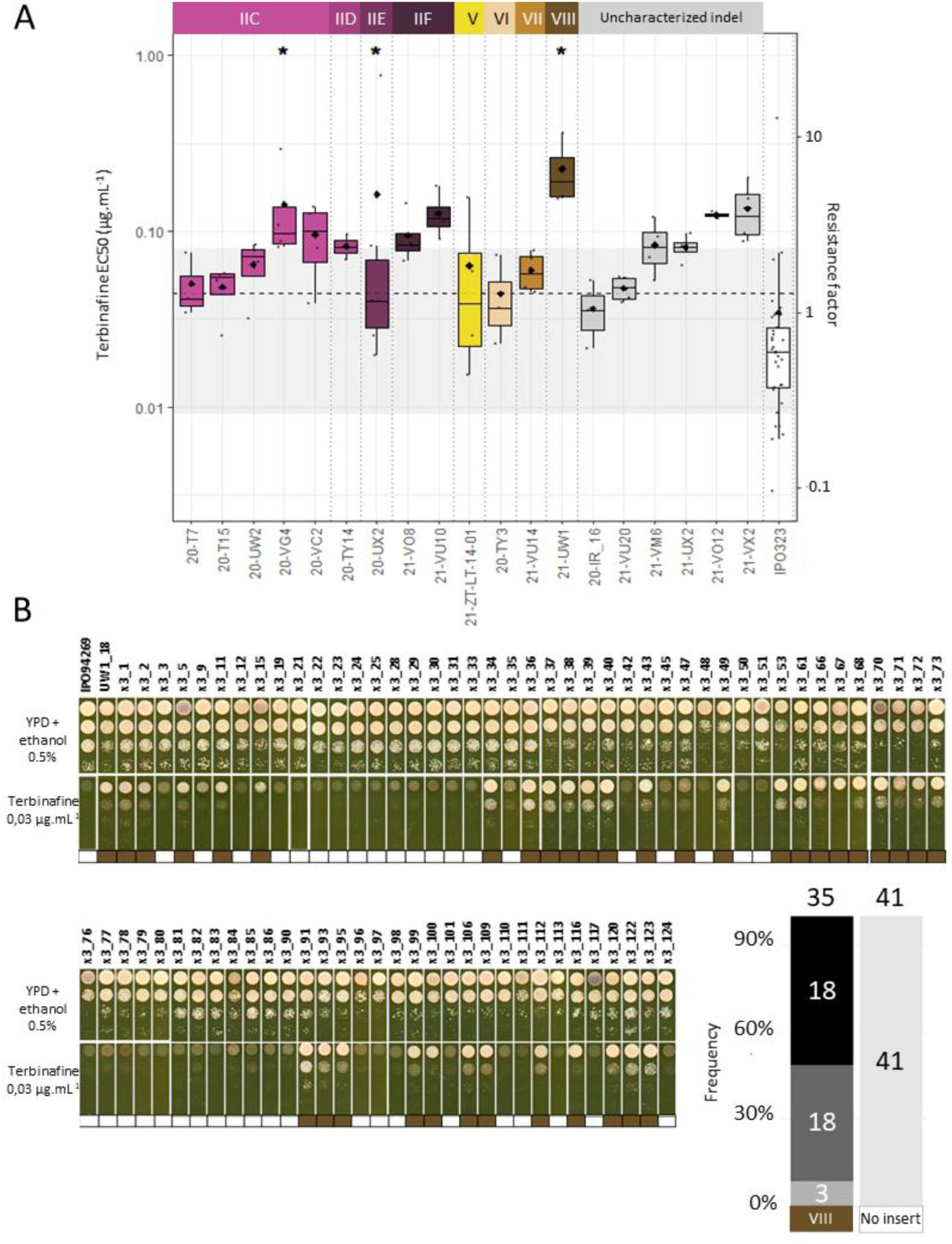
New P*MFS1* insertions differentially impact MDR phenotypes in *Z. tritici*. **A. Terbinafine *EC₅₀* of strains carrying new P*_MFS1_* insertion.** For EC₅₀ determination, growth in liquid YPD medium supplemented with increasing terbinafine concentrations was monitored at OD405_nm_ after 7 days (18°C, dark), and expressed as percentage of growth relative to control. EC₅₀ was computed from dose-response curves fitted to each technical replicate. Each strain was tested in at least two biological replicates (individual points in boxplots), each with three technical replicates. Statistical differences relative to the sensitive control strain IPO323 were tested on log-transformed values using a linear mixed-effects model including biological replicate variability as random effect, with estimated marginal means compared *via* contrast post-hoc tests and Bonferroni correction (* *P* < 0.05). Strains with uncharacterized indels (NA in panel A) or no-insert but with upstream insertions detected by whole-genome sequencing (20-IR_16, 21-VM6, 21-UX2) are included. All *EC₅₀*values are represented log10-transformed for clarity. **B. Validation of P ^typeVIII^ contribution to MDR by forward genetics.** Growth of parental strains (IPO942696, *P_MFS1_^no insert^*; 21-UW1, *P_MFS1_^typeVIII^*) and their individual progeny was tested on solvent and terbinafine-supplemented YPD medium. Colored cells indicate P*MFS1* genotype of each progeny determined by PCR amplicon size: white = no insert, brown = VIII indel. Frequencies of HR (black), MR (grey), LR (light grey), and S (lighter grey) phenotypes associated with each genotype are shown in the bottom-right graph

*P_MFS1_^type III^* was also consistently associated with terbinafine resistance. In the qualitative screen, 90% (19/21) of strains displayed an HR relative profile (Fig. 2B). EC₅₀ measurements revealed RF values ranging from 3 to 7, generally exceeding those of the Ref_III strain (RF 3.0), which served as the internal control for this genotype (Fig. 3A-5). This variability suggests additional background effects. To test this, we examined the terbinafine resistance phenotype of the progeny of a sexual cross between two *P_MFS1_^type III^* strains (20-LQ2 and 20-WK1, respectively LR and HR). The overall mean terbinafine EC_50_ of the progeny was approximately 0.03 µg.mL^-1^, with a substantial standard deviation of 0.016 µg.mL^-1^. This standard deviation reflects the underlying distribution of EC₅₀ values among the progeny which showed a continuous, quantitative pattern (Fig. 3B), with a broad-sense heritability (H²) of ∼40%. This indicates that, beyond the *P_MFS1_^type III^* insert, other genetic factors contribute to the observed resistance. These results support a polygenic model of MDR even among strains sharing the same insert.

**Fig. 5.**
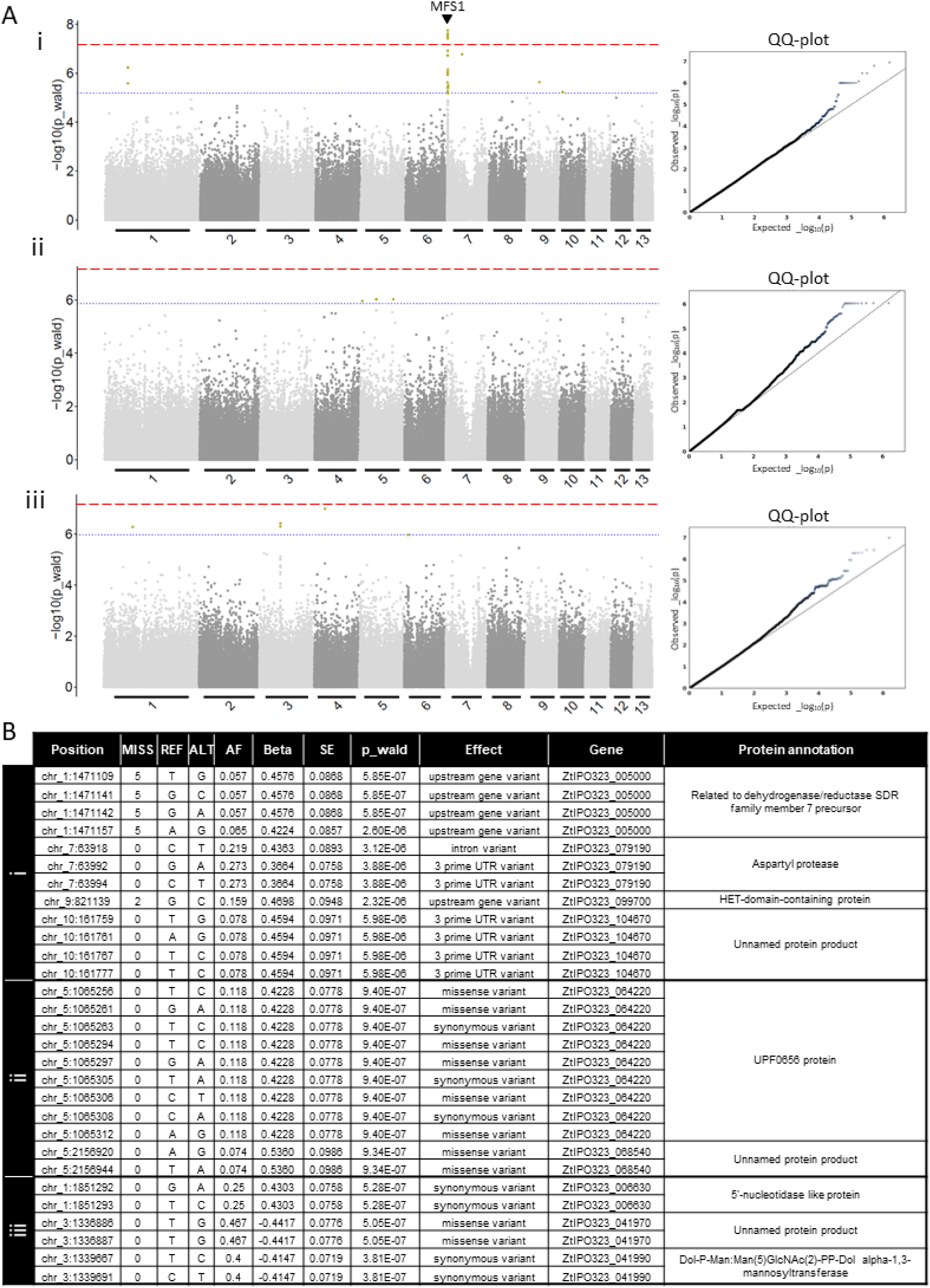
Genome-wide association confirms that *MFS1* remains the major locus associated with multidrug resistance (MDR), but additional loci may contribute to resistance variance. Genome-wide association tests for terbinafine resistance phenotypes. Manhattan plots (a, c, e) and Q–Q plots (b, d, f) are shown for each GWAS testing the association between SNPs and terbinafine resistance. Estimated marginal means of log-transformed EC₅₀ (Log_EC₅₀_) were extracted from a mixed linear model (*lmer*(Log_EC₅₀_ ∼ Souche + (1|Date))) and used as quantitative phenotypes for 128 strains (a, b) and corresponding subsets (c, e). Subsets were designed to mask the effect of the P*MFS1* locus through nested GWAS, performed separately on two groups of strains defined by the genotype of the top SNP from the complete GWAS: the reference allele group (c, 68 strains) and the alternative allele group (e, 60 strains). Complete, reference allele–nested, and alternative allele–nested GWAS analyses were performed using **GEMMA** on 758,800; 774,740; and 756,655 SNPs, respectively, fitting a linear mixed model (LMM). The first three principal components of the SNP-based PCA were included as covariates to account for population structure. On Manhattan plots, red and blue dashed lines represent Bonferroni and 10% FDR significance thresholds, respectively. SNPs surpassing these thresholds are highlighted in green. **B. Candidate SNPs for additional mechanisms contributing to MDR.** Candidate SNPs surpassing the 10% FDR threshold were extracted from each GWAS (a: complete dataset, b: reference-allele nested GWAS, c: alternative-allele nested GWAS). Only SNPs located outside the *MFS1* region and within 5 kb of annotated genes are shown. Columns indicate: **Missing:** number of strains with missing genotypes; **REF / ALT:** reference and alternative alleles; **Beta:** estimated effect size of the SNP on phenotype; **p_wald:** Wald test p-value for SNP–trait association; **Effect:** predicted impact of the SNP on gene expression or protein structure (predicted with SNPEff); **Gene:** nearest affected gene; **Protein annotation:** predicted protein function according to Lapalu *et al*., 2025.

The newly identified *P_MFS1_* variants showed variable MDR associations. Among the six strains harboring *P_MFS1_^type IIC^* tested for qualitative resistance, profiles ranged from sensitive (S) to HR (Fig. 2C). Only one of them exhibited a significantly high EC₅₀, suggesting an inconsistent link to resistance (Fig. 4A). The *P_MFS1_^type IID^* strain (20-TY14) displayed low resistance in qualitative screening and no significant EC₅₀ increase, indicating no effect of this insert on MDR. In contrast, the single *P_MFS1_^type IIE^* strain (20-UX2) showed high resistance qualitatively and a high EC₅₀, but with large variability across replicates. The *P_MFS1_^type IIF^* variant was observed in two strains with contrasting qualitative resistance profiles (one LR and one HR). However, EC₅₀ measurements showed no significant differences compared to the sensitive control, indicating that *P_MFS1_^type IIF^* is not consistently associated with increased terbinafine resistance. These results suggest that, unlike canonical *P_MFS1_^type II^* inserts (IIA and IIB), the newly identified *P_MFS1_^type II^* variants may not be sufficient on their own to confer MDR,or that additional epistatic genetic factors modulate its effect.

Strains with *P_MFS1_^type VI^*, *P_MFS1_^type VII^*, and *P_MFS1_^type V^* insertions also showed little or no association with terbinafine resistance. EC₅₀ measurements for these variants were not significantly different from the sensitive reference strain IPO323 (Fig. 4A). For *P_MFS1_^type VI^*, this lack of association was further confirmed by functional validation using a gene replacement approach. *P_MFS1_^type VI^* insert was introduced into IPO323 at the *MFS1 locus*, and the resulting transformants failed to grow on discriminatory concentrations of terbinafine or tolnaftate (Fig. S1). In contrast, the only strain carrying the new *P_MFS1_^type VIII^* variant (21-UW1) showed a HR phenotype in qualitative assay, further confirmed by a sevenfold increase of terbinafine EC_50_ compared to IPO323. Genetic validation through a sexual cross between 21-UW1 and the sensitive IPO94269 strain demonstrated complete co-segregation between the *P_MFS1_^typeVIII^* genotype and resistance among 79 progeny isolates (Fisher’s exact test, *p* < 2.2 × 10⁻¹⁶), confirming the causative role of this structural variant in MDR (Fig. 4B). Finally, strains genotyped as *P_MFS1_^no-insert^* displayed a broad range of resistance profiles, from sensitive to HR (Fig. 2B). EC₅₀ analysis identified two *P_MFS1_^no-insert^* strains with significantly higher EC_50_ mean compared to IPO323, including strain 21-VY4, with an RF of ∼3 (Fig. 3A-2). Additionally, none of the strains carrying uncharacterized variants detected with whole-genome sequencing showed significant increased EC₅₀ values compared to the reference strain (Fig. 4A), suggesting that these variants do not contribute to resistance under tested conditions. This indicates that resistance in some strains can arise independently of detectable promoter insertions, highlighting the possibility of distal or alternative genetic mechanisms contributing to MDR.

Altogether, our phenotyping results show that structural variants at the *P_MFS1_* promoter have differential impacts on MDR, in *Z. tritici*. While canonical insertions like types I, IIA, IIB, and III are strongly associated with MDR, other variants show limited or inconsistent effects, indicating that additional genetic factors may be involved in MDR.

### GWAS confirms *P_MFS1_* as the main contributor to MDR and supports polygenic determinisms of MDR in the field

To uncover the genetic determinants contributing to MDR in *Z. tritici*, we performed a genome-wide association study (GWAS) on 128 strains of our collection. As phenotype, we used the EC₅₀ values for terbinafine resistance (Fig. S2 and S3; Table S1).

In total, **68** SNPs were significantly associated with variation in terbinafine EC₅₀ values using a 10% false discovery rate (FDR) threshold (Fig. 5A, Table S2). The GWAS including all strains (Fig. 5A-i and B-i) revealed 49 significant SNPs in total. As expected, the most significant associations clustered at the telomeric end of chromosome 7, between positions chr7_22767 and chr7_36953, encompassing the *P_MFS1_* locus. Among these, the top-associated SNP (chr7_36375) is located within the *MFS1* coding sequence, and the second (chr7_35421) lies 12 bp upstream of the known 401 bp upstream insertion site. Both SNPs passed the Bonferroni-corrected threshold for significance and likely reflect linkage with structural variant polymorphisms at *P_MFS1,_* rather than direct causality. These results confirm *P_MFS1_* as a major resistance *locus* in the population. Beyond chromosome 7, several other *loci* were significantly associated with EC₅₀ values at 10% FDR. SNPs on chromosomes 1, 9, and 10 mapped near genes with potential roles in xenobiotic metabolism or transport (Fig. 5B-i). For example, 5 significant SNPs on chromosome 1 lie between *ZtIPO323_005000*, encoding a short-chain dehydrogenase/reductase (SDR), and *ZtIPO323_005010*, a predicted sugar transporter. The first one is plausible contributor to MDR, given known roles of SDRs in detoxification (De Rouck *et al*., 2023; Persson *et al*., 2009; Xue *et al*., 2020). Another significant SNP on chromosome 9 is close to *ZtIPO323_099690*, annotated as a homolog of the fluconazole resistance protein FLU1. Finally, we found 4 significantly associated SNPs on chromosome 10, in the 3’UTR of ZtIPO323_104670, a gene of unknown function. The GWAS results indicate that the *P_MFS1_ locus* explains the high proportion of phenotypic variance (15–21 %), while other *loci* also contribute substantially (14-17 %), indicating that terbinafine resistance is a quantitative trait influenced by multiple *loci*.

To uncover loci associated with MDR independently of *P_MFS1_*, we also performed two nested GWAS on subsets of strains carrying either the reference (REF; n = 68) or alternate (ALT; n = 60) allele at the top *P_MFS1_*-linked SNP (chr7_36375). The first one revealed 12 significant SNP and the second one 7. As expected, no significant SNPs were detected on chromosome 7 in either subset, confirming that the *P_MFS1_* signal was effectively masked. Several *loci* reached the 10% FDR threshold in these nested scans (Fig. 5A-ii,3, 5B-ii,iii), supporting the existence of additional MDR-related *loci* beyond the canonical *P_MFS1_* insertions. In the REF-GWAS, we identified 9 SNPs significantly associated (10% FDR threshold) overlapping the coding region of ZtIPO323_064220 encoding a tetratricopeptide repeat (TPR)-like protein and 2 significant SNPs overlapping ZtIPO323_068540, a gene of unknown function (Fig. 5A-ii, 5B-ii). The ALT-GWAS revealed three *loci* with 2 significant SNPs each, on chromosome 1, and 3, overlapping with ZtIPO323_00630 (nucleotidase-like protein), ZtIPO323_041970 (unknown function) and ZtIPO323_041990 (mannosyltransferase), respectively (Fig. 5A-iii, 5B-iii). These genes, despite no obvious association with non-target site resistance mechanisms, may still participate to a global regulation of MDR. Altogether, our GWAS results corroborate our phenotypic analyses, confirming that while the *P_MFS1_* locus is a major determinant of MDR in *Z. tritici*, multiple additional *loci* also contribute to resistance. This strengthens the conclusion that multidrug resistance behaves as a quantitative trait underpinned by a polygenic architecture.

## Discussion

In this study, we provide an extensive analysis of structural variation at the *MFS1* promoter in *Z. tritici* and reveal the complex spectrum of MDR genetic basis. By combining high-resolution phenotyping with forward and reverse genetics, as well as genome-wide association analyses across a large, recently sampled European population, we demonstrate that MDR is not only driven by major-effect transposable elements insertions (e.g., *P_MFS1_ types I, IIA, IIB,* and *III*), but is also shaped by a polygenic background. Our quantitative phenotyping confirms that even strains with identical *P_MFS1_* genotypes can differ markedly in resistance factors, and the GWAS further supports this view by identifying additional *loci* that may contribute to MDR. Together, our findings revise the current understanding of resistance evolution in *Z. tritici* by highlighting the interplay between transposon-driven structural changes and polygenic architecture in shaping MDR phenotypes in field populations.

The promotor region of *MFS1* is subject to diverse and recurrent transposon-mediated structural variation, in particular in this MDR targeted collection. We confirmed that nearly all the inserts described in previous studies (Mäe *et al*., 2020; Omrane *et al*., 2017, 2015) as well as the newly identified variants in the present work, exhibit derive from various TEs, including LTR/copia retrotransposon (type I), LINEs (type II), and MITEs (type VI and VII). These insertions are not randomly scattered across the promoter region but are concentrated within a narrow *locus*, often at shared or nearby insertion sites. Despite sharing insertion sites, phylogenetic analysis of flanking promoter regions suggests that most of the insertions arose independently. For instance, although all type II variants occupy the same insertion site 471 bp upstream of the *MFS1* start codon, they potentially derive from multiple, independent insertion events implying ongoing TE activity in the *P_MFS1_* region. These findings are consistent with recent work showing reduced TE silencing and active TE transcription in European *Z. tritici* populations (Abraham *et al*., 2024; Feurtey *et al*., 2023; Möller *et al*., 2021; Oggenfuss and Croll, 2023), particularly in subtelomeric and gene-rich regions such as the *MFS1* locus. Furthermore, we observed strong sequence conservation in the promoter sequences flanking insertions associated with MDR, specifically types I, IIA, IIB, and III, which suggests that these TE insertion polymorphisms (TIPs) may be under positive selection. Although the evolutionary forces shaping MDR-associated TIP remain unclear, TE insertions are known to contribute to standing genetic variation and may facilitate rapid adaptation under strong selection (Baril *et al*., 2025). Environmental stresses were shown to trigger TE de-repression (Fouché *et al*., 2020), and fungicide exposure may similarly act as both a selective pressure and as *stimulus* for TE mobilization, as proposed by Baril and Croll (2025). These findings underscore the dynamic interplay between TE activity, genome architecture, and adaptation in shaping the evolution of resistance in fungal pathogen populations.

Among the newly identified inserts in *P_MFS1_*, only the indel VIII was validated as conferring terbinafine resistance (and consequently MDR). This new *P_MFS1_* allele may increase in frequency due to its potential adaptive advantage.

Other newly identified TIPs were not associated with MDR under our experimental conditions. Since the isolates were initially selected based on their ability to germinate on terbinafine-supplemented media, the discrepancy could stem from differential expression of the resistance across developmental stages, morphotypes, or fungicide availability in the medium. These hypotheses may also explain inconsistencies between qualitative resistance scores obtained on solid media and EC₅₀ measurements in liquid culture. The latter, based on optical density, is strongly influenced by morphological transitions towards filamentous growth. In their study, Puccetti *et al*.(2025b, 2025a) found distinct genetic *loci* associated with fungicide resistance depending on the phenotyping method, highlighting the challenge of selecting the ecologically most relevant assay to describe resistance dynamics *in natura*. In addition, the germination of seemingly sensitive strains on terbinafine-supplemented media might reflect fungicide tolerance, an underrated concept in phytopathology despite its significant clinical implications (Berman and Krysan, 2020; Levin-Reisman *et al*., 2017).

Our results show that while the *P_MFS1_* insertion polymorphism is the major determinant of multidrug resistance (MDR) in *Z. tritici*, it is not the sole driver. We observed considerable variation in terbinafine resistance among strains carrying the same *P_MFS1_* genotype and in progeny from crosses between such strains. This indicates that MDR behaves as a quantitative trait shaped by additional genetic factors, which was confirmed by our GWAS analyses that identified candidate *loci* beyond *P_MFS1_*. These candidates include genes potentially involved in xenobiotic detoxification and membrane transport, consistent with resistance such as a short-chain dehydrogenase/reductase (SDR) and a homolog of the FLU1 transporter known to mediate MDR in *C. albicans* (Calabrese *et al*., 2000). Such polygenic architectures have been observed in other fungi such as *Botrytis cinerea* (Kretschmer *et al*., 2009; Leroux and Walker, 2013) and *Candida glabrata* (Huang *et al*., 2025). These findings are in alignment with those made by Puccetti and colleagues (Puccetti *et al*., 2025a) about the complex genetic nature of DMI resistance in *Z. tritici.* Moreover, while using terbinafine resistance is a convenient proxy for MDR large-scale screening, it may overlook a part of underlying genetic determinants. Additional *loci* associated with MDR but no affecting terbinafine resistance cannot be excluded.

The polygenic basis of fungicide resistance raises important implications for the evolution and management of resistance in pathogen populations. Even if individual genetic effects are small, cumulative resistance increases remain a major threat to treatment durability. This is especially critical for MDR, which compromises both current and future active ingredients, including biocontrol compounds, while enabling the survival of less susceptible isolates, possibly favoring the selection of resistance mechanisms with greater field impact. Our results highlight the urgent need for improved MDR prevention strategies. Indeed, while current recommendations promote the diversification of selection pressure through fungicide mixtures or alternations to delay target-site resistance (Rex Consortium, 2013), such approaches may paradoxically create selective conditions that promote multidrug resistance instead (Ballu *et al*., 2024, 2023, 2021; Barbosa *et al*., 2021; Comont *et al*., 2020; Lagator *et al*., 2013; Zhou *et al*., 2022, Neau *et al*., *in prep*). Thus, this work provides additional arguments in favor of limiting the use of AIs and diversifying selective pressures beyond chemical control within an integrated pest management framework (Rex consortium, 2016). By reducing the probability of recombination and co-accumulation of TSR and NTSR mechanisms in common genetic backgrounds, such strategies may help to preserve efficacy of phytopharmaceutical AIs.

## Experimental procedures

### Biological material

#### · Z. tritici 2020-2021 field strain collection

French field strains used in this study were obtained from the Performance Network (Garnault *et al*., 2019) during the years 2020 and 2021. Both French and British strains in our panel were isolated from infected wheat leaves and suspended in water following the protocol described by Garnault *et al*., (2019). The spore suspensions were plated on PDA media supplemented with 0.015 µg.mL^-1^ discriminatory concentration of terbinafine (Leroux and Walker, 2011) and incubated for 24 hours. After incubation, colonies with elongated germ tubes were picked using a sterile toothpick and isolated three to five times on PDA media supplemented with antibiotics (Kanamycin, Streptomycin, and Penicillin (Sigma-Aldrich) at 37.5 µg.mL^-1^). Strains from Belgium, Denmark, Estonia, Germany, Ireland, and Lithuania were provided by collaborators of the EURO-RES project (described in Hellin *et al*., 2021) as pure strains.

#### · Progeny from sexual crosses

Parental strains for crosses 21-UW1 x IPO94269 and 20-LQ2 x 20-WK1 were preliminary checked for mating type compatibility as described by (Waalwijk *et al*., 2002). Then, crosses were performed following the *in planta* procedure developed by Kema and collaborators (Kema *et al*., 1996) with adaptations described by Orellana-Torrejon and collaborators (Orellana-Torrejon *et al*., 2022). For the inoculation performed on June 6, 2023, blastospores of the parental strains were harvested from five-day-old solid precultures, resuspended in sterile water to a final volume of 20 mL and a concentration of 2.10^5^ spores.mL^-1^ (adjusted with a hemacytometer). The two suspensions of the parental strains were mixed at a 1:1 ratio, with final concentration of each strain reaching 1.10^5^ spores.mL^-1^ and a final volume of 40 mL. Two drops of the surfactant Tween 20 (Sigma, France) were added. These suspensions were sprayed with an atomiser (Ecospray, VWR, France) onto three adult plants (nine stems) of wheat Apache cultivar as described in Orellana-Torrejon *et al*. (2022). We promoted infection by enclosing plants immediately after inoculation for 72 h in a transparent polyethylene bag previously sprayed with distilled water. Plants were maintained in greenhouse environment for about three months, until complete drying (June to August 2023). Then, they were placed outdoor in August 2023 in Grignon (France, 78200) for five months in order to promote ascosporogenesis. In the residues of leaves and stems from plants of each cross were cut into 2 cm pieces dried at room conditions for one week. Progeny was harvested through experimental set up for ascospore ejection and catching. For this purpose, residues were soaked in water for 30 min and spread on dry filter paper in a crystal-clear polystyrene box (24 × 36 cm) leaving the lid half-open. Eight Petri dishes (90 mm in diameter) containing PDA medium were placed opened upside down 1 cm above the residues, for receiving the ascospores ejected from asci during residues drying process. Boxes were left at room conditions for 18 h before closing Petri dishes and incubating them under usual culture conditions. Between three to five days post-ejection, we regularly picked ascospore-derived colonies using sterile toothpicks and purified them through two successive single spore subcultures on PDA solid medium before long term storage. We harvested 76 individual progenies from 21-UW1 x IPO94269 and 151 from 20-LQ2 x 20-WK1 crosses, lately genotyped for their *P_MFS1_* insertion polymorphism as described hereafter.

All field strains, progenies and reference strains used in this study are listed in Table S1.

### General growth conditions

*Z. tritici* strains were preserved at −80°C in 25 % glycerol suspensions and precultured on solid YPD medium (20 g.L^-1^ dextrose, 20 g.L^-1^ peptone, 10 g.L^-1^ yeast extract, 10 g.L^-1^ agar) in the dark, 18°C, 60% humidity. For resistance phenotyping, strains were grown in 10 mL liquid YPD medium in 50 mL sterilized Erlenmeyer flasks plugged with cotton wool for 7 days at 150 rpm. Cell concentrations were determined by OD measurements at λ=405 nm in a Spectramax M2 microtiterplate reader (Molecular Devices, USA) according to the formula established by Ballu (Ballu, 2021).

### Fungicide preparation

Terbinafine (Sandoz SA, BALE/Switzerland) was solubilized in either DMSO or 80% ethanol (v/v). A concentrated solution of 4,000 µg.mL^-1^ was prepared and stored at 4°C. To prepare the fungicide-supplemented media, serial dilutions were performed in solvent, and the concentration was adjusted for the amount of solvent introduced to not exceed 0.5% of the final volume, in order to prevent solvent-induced toxicity (Ballu *et al*., 2021, 2022, 2023).

### Qualitative resistance assay to fungicides (droplet growth test)

The qualitative assessment of terbinafine resistance as proxy for the MDR phenotype was performed using a droplet test according to Ballu *et al*. (2021). Quickly, serial water dilutions containing 10^7^, 10^6^, 10^5^, and 10^4^ spores.mL^-1^ were deposited as approximately 5 µL drops onto square Petri dishes (12 cm) containing solid YPD medium supplemented with antibiotics (Kanamycin, Streptomycin, and Penicillin (Sigma-Aldrich) at 37.5 µg.mL^-1^), and a previously determined discriminatory concentration of fungicide) or solvent alone (see fungicide preparation). The discriminatory dose of terbinafine (0.03 µg.ml^-1^) was optimized on characterized pure strains to allow full growth of HR strains while preventing growth of the reference strain IPO323 (S strain). Fungal growth was scored after 7 days of culture. Each strain was scored with a number representing growth intensity, ranging from 0 (no growth) to 4 (growth at the lowest spore concentration). According to this score, strains were categorized as highly resistant (3 to 4, HR), moderately resistant (2, MR), low resistant (1, LR), or sensitive (0, S). Specifically for terbinafine resistance screening in field strain collection and 21-UW1 x IPO94269 progeny, Fisher’s exact test was used to assess the association between indel VIII and terbinafine resistance (sensitive versus resistant, with low, moderate, and high resistance grouped together) among the 21 UW1 × IPO94269 progenies (Table S3).

### Quantitative terbinafine resistance assessment ( EC₅₀ determination through dose response curve)

We performed quantitative terbinafine resistance assessment for a 126-strain subset of the 2020-2021 strain collection and 151 progenies from 20-LQ2 x 20-WK1 cross (Table S4). To the first set, we added four *P_MFS1_^no-insert^* field strains obtained through the Performance network between 2012 and 2013 (Table S1, Garnault *et al.,* 2019), sensitive to terbinafine (not germinating on PDA supplemented with 0.015 µg.mL^-1^) for establishing the basal field terbinafine EC₅₀.

We computed the EC₅₀ of terbinafine, corresponding to the concentration inducing its half-maximum inhibitory effect on fungal growth. For doing so, the spore concentration of the 7-day-old liquid precultures was adjusted to 1.25 × 10^5^ spores.mL^-1^ in 96-well microplates (Sarstedt) in a final volume of 200 µL of YPD medium supplemented with antibiotics (37.5 µg.mL^-1^ Kanamycin and Streptomycin (Sigma-Aldrich)) and terbinafine at a range of different concentrations or solvent alone (control). Different terbinafine ranges were tested for high resistance (HR) and low resistance (LR) strains (Table S5). A breathable film (Breathe-easy®; Diversified Biotech) was applied on microplates in order to limit evaporation and contaminations before starting a 7-day incubation at 18°C and 150 rpm in a microplate shaker (Innova S44i, Eppendorf). Each strain was tested in three technical replicates (three wells per fungicide condition in the same plate) and at least 2 independent biological replicates (independent days). As internal control, we systematically add IPO323 reference strain as negative control to every biological replicate. As positive controls for the 2020-2021 MDR collection we tested as well IPO323 isogenic transformants carrying either the *P_MFS1_*-type I allele, *P_MFS1_*-type II allele, or *P_MFS1_*-Type III allele integrated at the *MFS1 locus* of IPO323 (Omrane *et al*., 2017), renamed Ref-I, Ref-II, and Ref-III, respectively (Table S1). Among biological replicates including progenies from 20-LQ2 x 20-WK1 cross, we also systematically tested both parental strains.

After 7 days of growth, spore concentrations were determined from OD405 nm values (Ballu *et al*., 2021) and expressed as percent of growth of the control condition (solvent alone). EC₅₀ were computed from dose-response curves fitted to each technical replicate, using the R software (R Core Team, 2025) and drc package. After filtering data for quality, several dose–response models (LL.5, LL.4, LL.3, LL.2, W2.2, and W1.2) were fitted and the model with the lowest Bayesian Information Criterion (BIC) value was selected. Log-transformed EC₅₀ mean per strain and biological replicate (mean of technical replicates) were modeled using a linear mixed-effects model (lmer function from lme4 package (Bates *et al*., 2015), with “strain” as a fixed effect and “biological replicate” (experiment date) as random one. Model assumptions were verified using Levene tests for homoscedasticity and residual inspection, QQ-plot, and Kolmogorov–Smirnov test on standardized residuals for normality assumption (Fig. S2). We calculated estimated marginal means (emmeans package (Lenth, 2025)) for each strain, and performed statistical comparisons with contrast function. P-values were adjusted for multiple testing using the Bonferroni correction for each set of comparisons and are presented in Table S1. Raw data for both MDR field strains and 20-LQ2 x 20-WK1 progeny are available (Tables S6 and S7).

Broad-sense heritability (H²) of log-transformed terbinafine EC₅₀ among progenies of the 20-LQ2 x 20-WK1 cross was estimated fitting a linear mixed model including Strain and Date as random effects as: 𝐿𝑜𝑔(𝐸𝐶50)_𝑖𝑗𝑘_ = 𝜇 + 𝑆_𝑖_ + 𝐷_𝑗_ + 𝜀_𝑖𝑘𝑗_with μ the overall mean (the intercept), S_i_ the random effect of strain, D_j_ the random effect of experimental date and the residual error. H² was calculated as the proportion of total phenotypic variance explained by the strain effect as 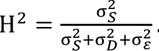.

### *P_MFS 1_* PCR-based genotyping

For DNA extraction, 50 – 100 mg of fresh spores were harvested per strain from 7-day old cultures on YPD agar media, placed in a microtiter plate and immediately frozen at −80°C. Cell lysis was performed mechanically twice in AP1 buffer supplemented with Rnase A and reagent DX (Qiagen) using tungsten beads in MIXER MILL MM 400 homogenizer (RETSCH®) at 20 Hz for 20 seconds. DNA was precipitated with sodium acetate (0.5 M final concentration, pH 5) and the addition of 0.7 vol of isopropanol. After centrifugation, dry DNA was resuspended in AE buffer (Qiagen). Genotyping of the *MFS1* promotor was performed on all 398 strains by PCR using the primer pair *P_MFS1_*_2F and *P_MFS1_*_4R and PCR condition described by (Omrane *et al*., 2017). All amplicon displaying length compatible with *P_MFS1_*-Type II and *P_MFS1_*-Type III alleles were sequenced with the same primers using Sanger protocol (Eurofins Genomics, Köln, Germany). Among others, 21 isolates with *P_MFS1_*-type I and 16 isolates with *P_MFS1_*-no insert length compatible amplification were also sequenced. Facing amplification failures for some strains (Table S1) we designed another pair of primers, ZtIPO323_079130-3F and ZtIPO323_079140-bR in coding sequences of both *MFS1* (ZtIPO323_079130) gene and upstream gene ZtIPO323_079130 (Table S8). We performed this last PCR using Taq polymerase Phusion® (Thermo Fisher Scientific Inc., Waltham, MA, USA) and adapted PCR conditions (Table S8) before sequencing with the same primers using Sanger protocol (Eurofins Genomics, Köln, Germany).

### Whole genome sequencing and variant calling

Genomic DNA was isolated from fresh fungal material collected from 7-day-old cultures grown on YPD agar (one 60 mm diameter Petri dish) for each of the 132 sequenced isolates (Table S1). DNA extraction was carried out using the DNeasy Plant Mini Kit (Qiagen) following the manufacturer’s protocol. DNA integrity was verified by agarose gel electrophoresis, and its concentration and purity were assessed using a NanoDrop spectrophotometer (Thermo Fisher Scientific). Extracted DNA was stored at –20 °C until sequencing.

Paired-end Illumina sequencing (150 bp, 25× coverage) was performed by BMKgene (Biomarker Technologies GmbH, Germany). Adapter sequences and low-quality regions were removed from raw reads with Trimmomatic v0.39 (Bolger *et al*., 2014), applying leading and sliding window trimming at Q ≥ 28 and discarding reads shorter than 50 bp. The cleaned paired reads were aligned to the Zymoseptoria tritici IPO323 reference genome (Goodwin *et al*., 2011) using BWA-MEM v0.7.17 (Li and Durbin, 2009) and resulting SAM files were converted to sorted and indexed BAM files with samtools v1.20 (Danecek *et al*., 2021). PCR duplicates were identified and removed using Picard v2.21.6 (Broad Institute, 2019).

Variant discovery was performed for each sample using GATK v4.4.0.0 HaplotypeCaller in GVCF mode (ploidy = 1) (Poplin *et al*., 2017), and individual GVCFs were combined into a single dataset. Variants were filtered with GATK VariantFiltration using the following thresholds: FS > 10, MQ < 20, QD < 20, and DP < 3. Additional filters excluded variants with ReadPosRankSum, MQRankSum, or BaseQRankSum values outside [–2, 2], to remove potential biases in mapping position, base quality, or alignment. Functional annotation of variants was performed using SnpEff v5.1 (Cingolani *et al*., 2012) with a custom database built for Z. tritici IPO323. For downstream analyses, the variant effect with the highest predicted impact (according to the SNPEFF effect score) was extracted using a custom Python script. Variant positions were then intersected with gene annotations from (Lapalu *et al*., 2025).

Whole-genome sequencing data generated in this study have been deposited in the NCBI Sequence Read Archive (SRA) under BioProject accession number PRJNA1444912.

### Genome Wide Association analysis and population structure

Prior to GWAS analysis, per-sample sequencing quality metrics were computed using VCFtools (v0.1.17) (Danecek *et al*., 2011) to estimate both the proportion of missing genotypes per individual and the mean sequencing depth across core chromosomes. Isolates were filtered based on missing data and coverage thresholds: individuals showing ≥20% missing genotypes or an average sequencing depth ≤6X were excluded from downstream GWAS analyses (Fig S2). Relatedness between isolated was assessed using a square identity-by-state matrix computed with PLINK (Chang *et al*., 2015) to identify clonal strains (Fig. S4).

As tested phenotype, we input emmeans of Log(EC₅₀) extracted from mixed linear model described in previous sections fitted on a set of strains excluding ones with >40% deviation between predicted and observed values (Fig. S2, 20-UX2 and 20-UX5). The final set of 128 strains tested in GWAS and associated emmeans is presented in Table S1.

Association tests were performed with GEMMA (v0.98.3; (Zhou and Stephens, 2012)) included in vcf2GWAS wrapper (Vogt *et al*., 2022). Genotypes were provided as a vcf input filtered as described before, including all chromosomes. A linear mixed model (LMM) was fitted, adding the first three principal components as fixed-effect covariates to account for population structure, setting parameters --cfile PCA and --covar 3 in vcf2gwas function. A kinship matrix, calculated from the genotype matrix by vcf2gwas, was automatically included as a random effect in the LMM to correct for relatedness among isolates. 758,800 biallelic SNPs with a minor allele frequency ≥0.05 were retained through vcf2GWAS per-preprocess as input for GEMMA. Nested GWAS were performed on two subsets of strains carrying either the reference or alternative allele at the most significant SNP from the initial GWAS (68 and 60 strains, respectively), with 774,740 and 756,655 SNPs retained for analysis. SNPs with p-value lower than FDR <10 % were considered candidate variants (Table S2). Percent variance explained by each SNP was computed as in (Kumar *et al*., 2021) (Table S2).

We reproduced principal component analysis (PCA) upon 14806 retained biallelic SNPs (100 % call rate, minor allele frequency ≥0.05, thinned to one variant per 1 kb, Linkage desequilibrium-pruned markers (r² < 0.2)), either including all strains or after retaining a single representative per clonal group. The first two principal components were visualized in R and provided in Fig. S4.

### Sequence analysis

For sequence analysis of proximal and distal *P_MFS1_* promoter, we combined sequences obtained by Sanger sequencing of PCR amplicons, and sequences obtained from WGS illumina sequences reassembled genomes (described in previous *P_MFS1_* genotyping and Whole genome sequencing sections). Individual genome assemblies were generated using SPAdes v3.15.5 (Bankevich *et al*., 2012) from paired-end reads (R1 and R2) for each isolate. Sequences were designed as hooks (conserved sequences in all isolates) in CDS of ZtIPO323_079130 (5’ hook, Chr7_33925-33950 “GAGAGGAGAATTCGCTGAGGAGGGAT”) and *MFS1* (3’ hook, Chr7_37216-37259 “CAACCTGGCGCTCCAAACTCCAACAATTCGACATCTTCGGCACC”) and used to locate and extract *P_MFS1_* from each assembly (10 kb downstream to 5’ and upstream of 3’ hooks) using seqkit (Shen *et al*., 2016) and SAMtools (Danecek *et al*., 2021). Sequences extracted with both hooks were aligned with Kalign algorithm (Lassmann, 2020) and manually curated for obtaining full *locus*. Because repetitive sequences of inserts can compromise assembly from short reads, Sanger sequences were integrated to confirm and complete *P_MFS1_* sequences for defining a consensus sequence per isolate. When insertion polymorphisms were detected only in the reassembled genome upstream of the Sanger-genotyped region, they were retained but flagged as potentially unreliable. Only their insertion sites within the promoter were considered informative. Multiple sequence alignments (MSAs) of all *P_MFS1_* sequences were generated with MUSCLE (Edgar, 2004) and manually adjusted (see data availability).

For investigation of insertion origin, known and novel insert sequences were blasted against the annotated IPO323 reference genome annotated for transposable elements (Baril and Croll, 2023). Conservation of the insert sequences among recent field isolates was assessed *via* blastn against *Z. tritici* NCBI experimental databases.

MSA of *P_MFS1_* was transformed into VCF format with custom Python scripts, capturing variant positions and indels. Haplotypes were defined as sequences with distinct combinations of indels >100 bp, and these haplotypes were used to generate synteny maps with gggenome in R (Hackl *et al*., 2024). The *P_MFS1_* MSA filtered for positions present in all isolates (2074 positions) was used for phylogenetic analysis of conserved regions. Consensus tree was inferred with IQ-TREE v2.2.2.6 using the HKY+F+I+R2 model (Minh *et al*., 2020). For whole-genome phylogeny, MSA were generated from previously described whole-genome VCFs. Variants were filtered to retain biallelic SNPs with minor allele frequency >5%, thinned to one SNP per 1 kb, and LD pruning was performed with a 50-SNP sliding window moving 5 SNPs at a time at r² > 0.2 using PLINK. Remaining 14,833 variants were converted to PHYLIP format with vcf2phylip.py (Ortiz, 2019) and used to build consensus trees in IQ-TREE (GTR+F+ASC+R5 model). For both analysis, branch support was assessed with 1,000 ultrafast bootstrap replicates, and trees were visualized with iTOL v7 (Letunic and Bork, 2024). Nucleotide diversity (π) was computed with pegas in R (Paradis, 2010) from both *P_MFS1_* MSA and the whole-genome SNP set, among strains sharing the same *P_MFS1_* insertion polymorphism haplotype using only one representative per clonal group.

### *P_MFS1_^typeVI^* gene replacement strategy

To introduce the *P_MFS1_^typeVI^* allele into the sensitive IPO323 strain, a replacement cassette was built using the same strategy described in Omrane *et al*. (2017) with minor changes. For all PCR performed to obtain cloning fragments, the Phusion® Taq polymerase (Thermo Fisher Scientific Inc., Waltham, MA, USA) was used under adapted PCR conditions using primers referenced in Table S8. The region ranging from 1,273 bp upstream of *MFS1* until 510 bp downstream of the open reading frame (ORF), was amplified from 20-TY3 genomic DNA with the primer pair MDR6-7_pKr_F/ Gibs_MDRIV_R. A 737-bp 3’ flank of the *MFS1* gene to facilitate homologous recombination was amplified from IPO323 genomic DNA with primers Ipo323_hyg_F and Ipo323_pKr_R. Finally, the hygromycin resistance marker gene *hph* was amplified from plasmid pCAMB-HPT-Hind (Kramer *et al*., 2009) with the primer pair Gibs_Hygro_for/Hygro_ipo323_R (Table S8). The three fragments were assembled with XhoI-EcoRI-digested pCAMB-HPT-Hind using the Gibson Assembly Cloning kit (New England Biolabs, Ipswich, MA, USA) according to the supplier’s instructions. NEB 5-alpha competent *E. coli* (New England Biolabs, Ipswich, MA, USA) were transformed by heat shock with 2 µl of the assembly reaction mixture. Successfully transformed colonies were selected on medium added with kanamycin (50 mg.L^-1^) then validated by PCR on colonies with 3 pairs of primers pCAMBIA_rev/*MFS1*_rev, *MFS1*_for/pHygro_rev and pHygro_for/pCAMBIA_for (Table S8). Positive clones were picked for plasmid extractions and mini-prepped plasmid construct was validated by Sanger sequencing (Eurofins, Luxembourg) with primers listed in Table S8.

*Agrobacterium tumefaciens* strain AGL1 was transformed by heat shock with the generated plasmid pCAMBIA0380_RH_*P_MFS1_*_V_*MFS1*_hygro. Positive colonies were selected on YEB broth (beef extract 5 g.L^-1^, yeast extract 1 g.L^-1^, peptone 5 g.L^-1^, saccharose 5 g.L^-1^, MgCl2 0,5 g.L^-1^) with rifampicin (50 mg.L^-1^), kanamycin (50 mg.L^-1^), and ampicillin (100 mg.L^-1^) then screened by same colony PCR as for *E.coli*.

The IPO323 reference strain was transformed *via A. tumefaciens*-mediated transformation (ATMT) following a procedure adapted from Bowler *et al*. (2010). An *A. tumefaciens* strains carrying the pCAMBIA0380_RH_*P_MFS1_*_V_*MFS1*_hygro plasmid was grown overnight at 28°C, 200 rpm in liquid YEB medium with antibiotics as described before. Cells were diluted in induction medium (IM, recipe table S9) containing 50 µg.mL^-1^ kanamycin and 40 µg.mL^-1^ acetosyringone, to a final OD_660nm_ of 0.4 and mixed v/v with 10^7^ spores.mL^-1^ IM suspension of IPO323 blastospores harvested from 7-day solid YPD cultures. Mixtures were plated onto IM-agar plates covered with cellophane membranes (Cellophane Membrane Backing 165-0963, Biorad) and incubated for 48h at 18 °C before transferred onto MM-Zt (recipe table S9) plates supplemented with 250 µg.mL^-1^ cefotaxime (Kalys SA) 100 µg.mL^-1^ Streptomycin (Sigma), 50 µg.mL^-1^ Spectinomycin (Fluka) and 200 µg.mL^-1^ Hygromycin (Sigma)for 15 to 20 days at 18 °C. Transformants were isolated and selected on MM-Zt plates containing hygromycin (200mg.L-1) for 10 days at 18°C. before purification by single colony propagation twice. Successful allele replacement at the locus was confirmed by colony PCR (primers listed in Supplementary Table S8).

## Data availability

Supplementary figures, tables, and plasmid maps used in this study, as well as raw data and analysis scripts are publicly available on Data.gouv.fr under the DOI: https://doi.org/10.57745/NGQAXE.

## Supporting information

plasmid map

## Acknowledgements

SPL was supported by Anova-Plus Company. EN had a travel grant by University Paris-Saclay, Biosphera Graduate school and by INRAE division “Plant Health & Environment”. Cécile Lorrain is funded by an SNSF Ambizione grant (PZ00P3_209022). This work was partially funded by the Center for Interdisciplinary Studies on Biodiversity, Agroecology, Society, and Climate (C-BASC) of the Paris-Saclay University. The authors thank the Performance network as well as participants of the EURO-RES project (C-IPM ERANET) for providing leaf samples or isolates in 2020–2021. The authors are grateful to Frédéric Suffert and Manon Delanoue for assistance with crosses of *Zymoseptoria tritici*, to Anne Genissel for guidance on GWAS analyses, and to Anne-Lise Boixel for precious help with statistical analyses. BIOGER benefits from the support of Saclay Plant Sciences-SPS (ANR-17-EUR-0007).

## Supplementary figures

**Fig. S1:**
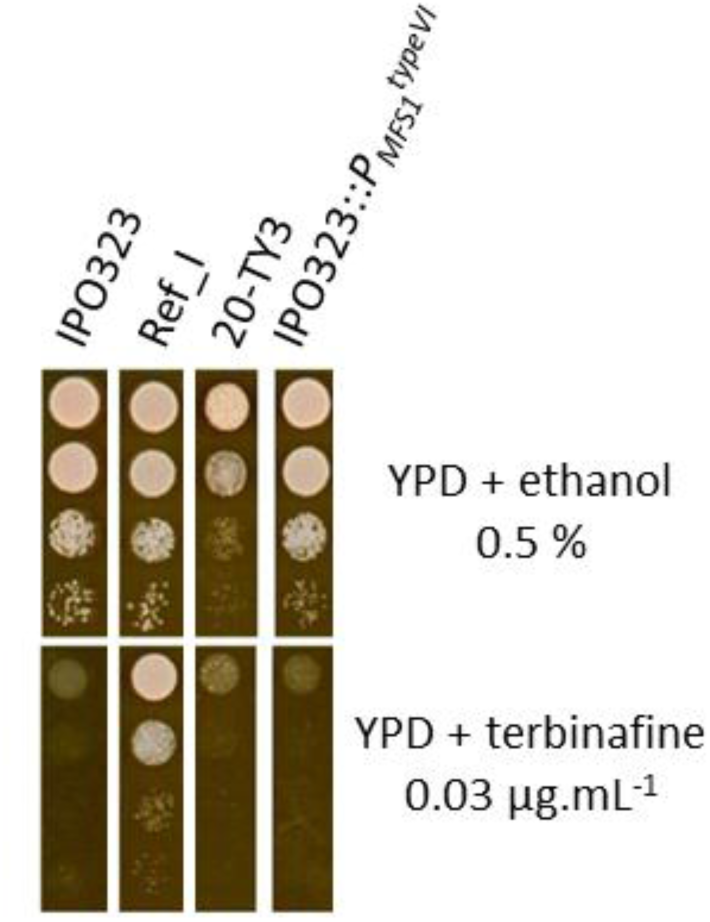
PMFS1typeVI transposon insertion is not sufficient to induce terbinafine resistance in the IPO323 genetic background. For these droplet tests, IPO323:*P_MFS1_^typeVI^* mutant, natural *P_MFS1_^typeVI^* strain 20-TY3, susceptible control IPO323 and positive MDR control Ref_I were precultured for 7 days and approximately 5 µL of serial spore dilutions (10^7^, 10^6^, 10^5^ and 10^4^ spores.mL^-1^) were spotted onto solvent or terbinafine-supplemented YPD medium. Growth was assessed after 7-day culture. IPO323:*P_MFS1_^typeVI^* appears as susceptible as IPO323 control strain.

**Fig. S2:**
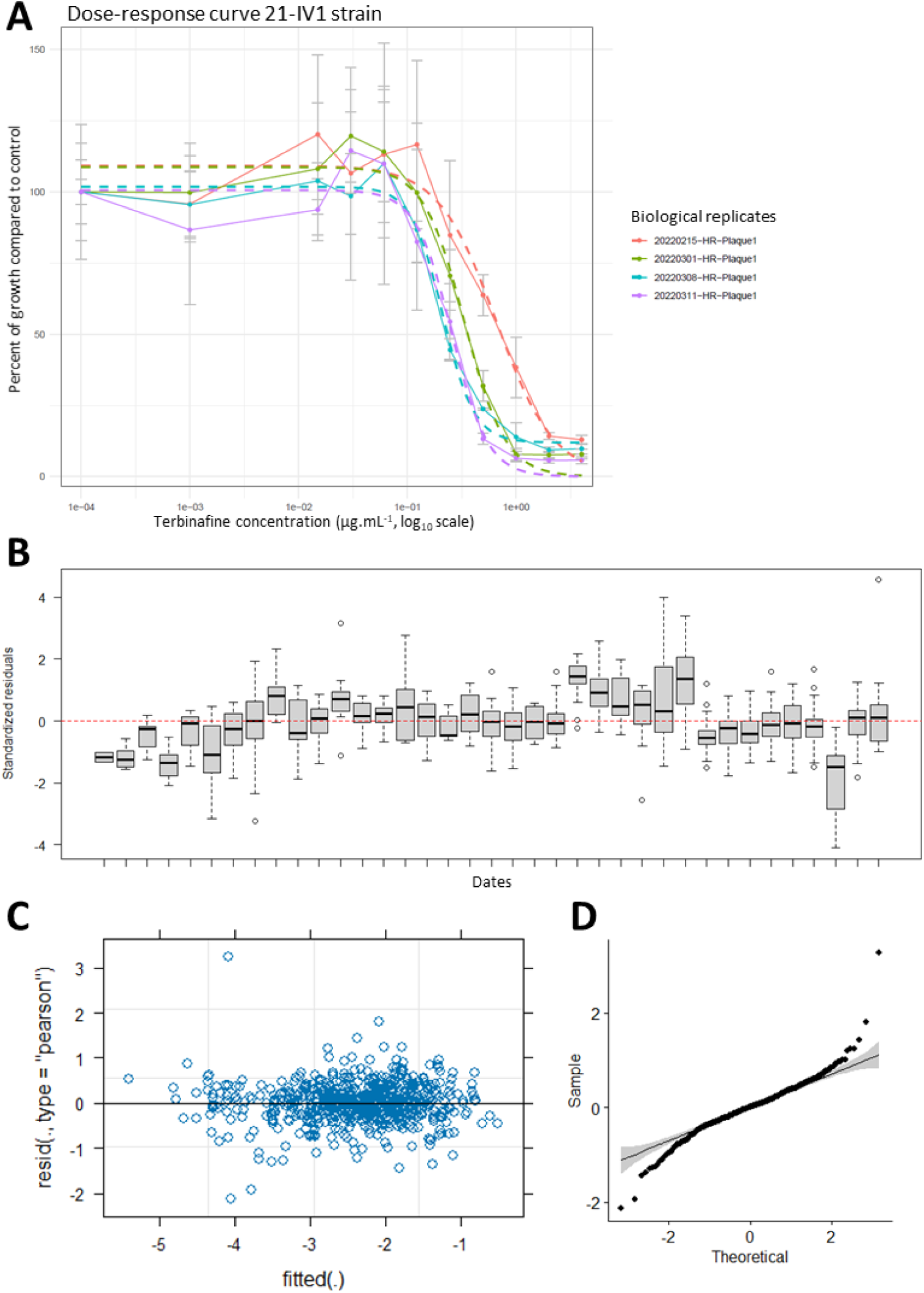
Assessment of experimental variability and statistical modeling of terbinafine EC₅₀ data. A: Example of terbinafine dose–response curves obtained for the 21-IV1 strain. Each curve was fitted for one biological replicate based on three technical replicates per terbinafine concentration (error bars represent the standard deviation of technical replicates). B: Standardized residuals from the linear model LogEC₅₀ ∼ Strain plotted for each biological replicate (experimental date), highlighting substantial experimental variability. This observation justified the inclusion of Date as a random effect in the mixed linear model used for statistical analyses. C and D: Diagnostic plots of the mixed linear model used for statistical comparisons of terbinafine EC₅₀ in field strains, including experimental date as a random effect (LogEC₅₀ ∼ Strain + (1|Date)). The distribution of Pearson standardized residuals versus fitted values (C) and the QQ plot (D) indicate acceptable homoscedasticity and approximate normality of residuals.

**Fig. S3:**
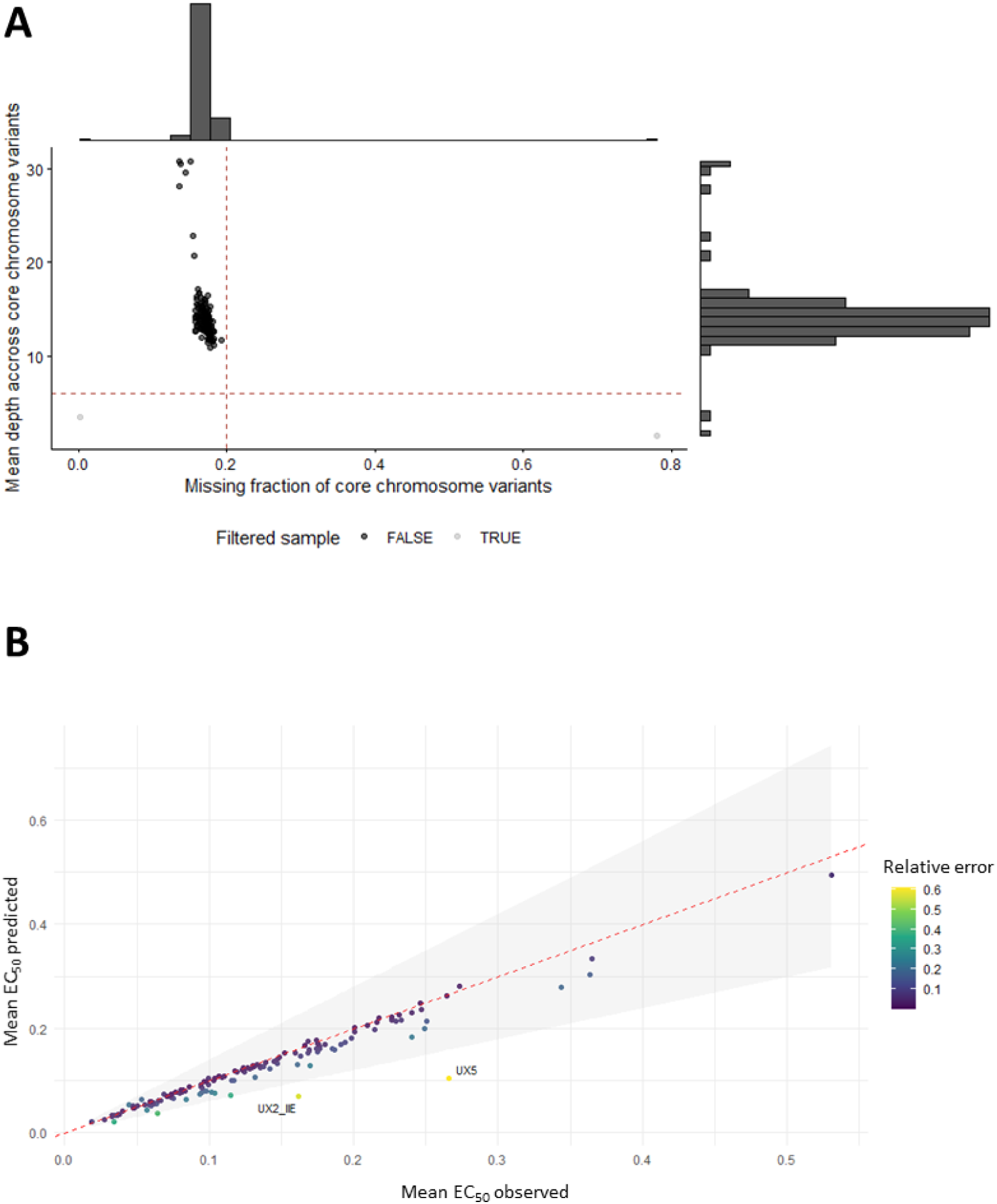
Sample quality control for GWAS. **A: Quality control based on missing data and sequencing depth.** Each point represents one strain, plotted according to the fraction of missing genotypes (F_MISS) and the mean sequencing depth across core chromosome variants. Dashed red lines indicate the filtering thresholds used for GWAS analyses (F_MISS ≥ 0.2 and mean depth ≤ 6). Strains failing at least one of these criteria are shown in grey, whereas retained strains are shown in black. Marginal histograms display the distribution of missing data and sequencing depth across strains. This filtering step was applied to exclude low-quality samples prior to downstream GWAS analyses. **B: Quality control based on quality of mean EC_50_ prediction.** Observed versus predicted mean EC₅₀ values colored according to relative prediction error. The shaded grey area represents a ±40% deviation from the identity line. Strains exceeding this threshold (UX2-IIE and UX5) were considered poorly predicted by the model and excluded from downstream GWAS analyses.

**Fig. S4:**
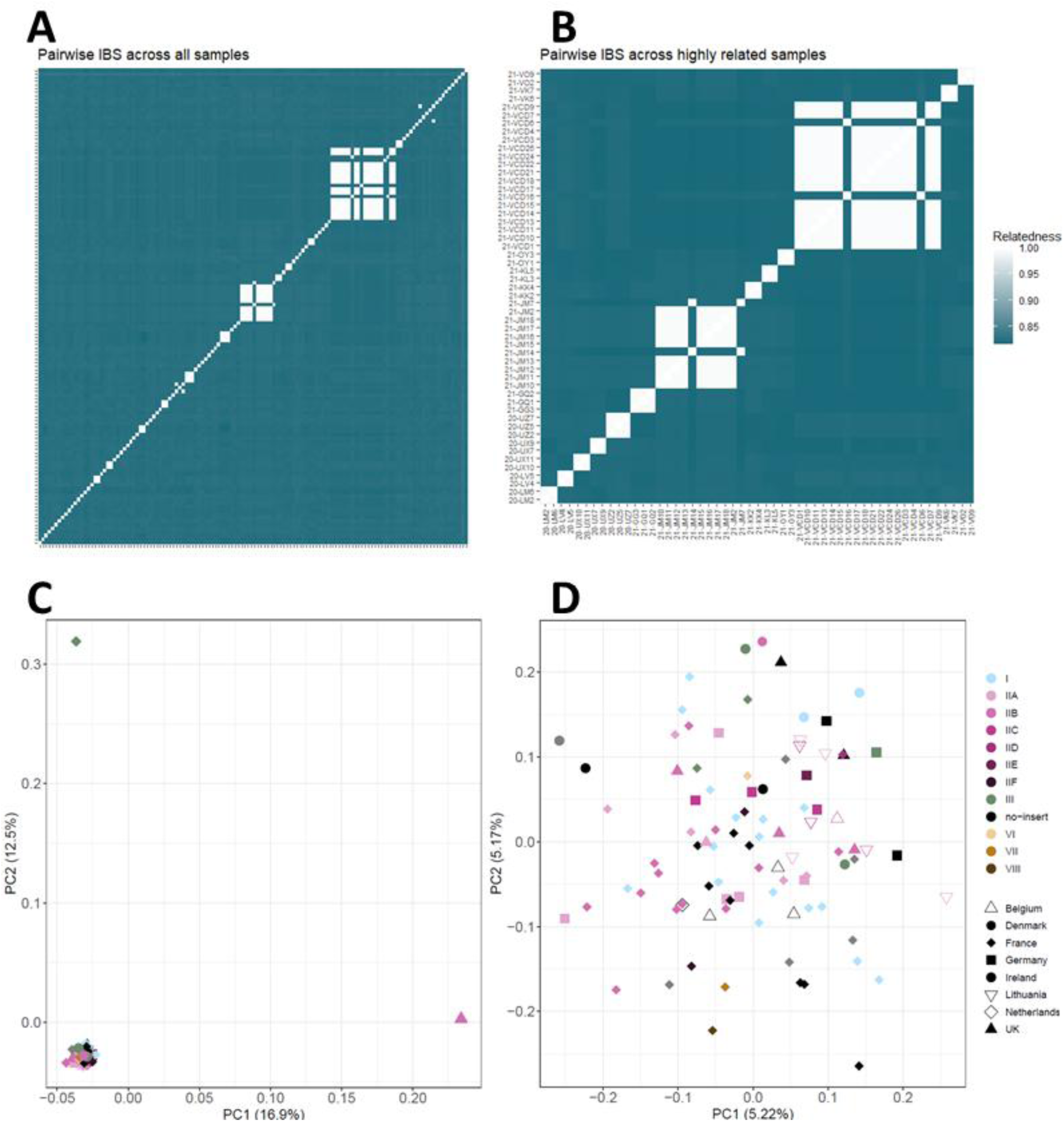
Kinship and population structure analyses. **(A, B) Pairwise genetic relatedness among MDR field strains.** (A) Heatmap of pairwise identity-by-state (IBS) values computed from genome-wide SNP data for all strains retained after quality filtering. lighter colors indicate higher genetic relatedness. (B) Heatmap restricted to strains involved in highly related pairs (IBS ≥ 0.99). This representation highlights clusters of near-clonal or highly related strains. **(C, D) Principal component analysis (PCA) of population structure** performed on LD-pruned, biallelic SNPs (MAF ≥ 5%, no missing data, thinned to one SNP per 1 kb). Each point represents a strain, colored according to P*MFS1* genotype and shaped by country of origin. (C) PCA including clonal strains. The presence of tight clusters reflects the contribution of clonal or near-clonal strains to population structure, motivating the inclusion of the first three principal components as covariates in GWAS analyses. (D) PCA after random selection of a single representative per clonal cluster reveals that population structure is neither driven by P*MFS1* insertion type nor by geographic origin.

